# Scaling relations of multicomponent phase coexistence boundaries

**DOI:** 10.1101/2025.04.02.645070

**Authors:** Daoyuan Qian, Julia Acker, Rob M. Scrutton, Charlotte M. Fischer, Seema Qamar, Tomas Sneideris, Alex Borodavka, Tuomas P. J. Knowles

## Abstract

Multicomponent phase-separating molecular systems play a key role as membraneless organelles in living cells. Phase diagrams are indispensable for probing the concentration-dependent behaviour of these condensates, yet their interpretation has remained largely qualitative due to the challenges of modelling complex, multicomponent interactions. Here, we present a generic framework for quantitative analysis of phase diagrams. We derive an analytical expression for the phase boundary normal vector, which reveals two distinct scaling regimes: one for associative phase separation, characterised by a power-law scaling analogous to mass-action kinetics that allows for direct extraction of dense phase solute molar ratios, and another for condensate dissolution, where an exponential scaling quantifies three-body repulsion effects triggered by the addition of a new component. We demonstrate the practical utility of our framework by applying it to a range of experimentally measured phase diagrams, including those for proteins such as NSP5, NSP2, FUS, Whi3, G3BP1, and antimicrobial peptides. Collectively, our work not only provides a clear quantitative interpretation of phase diagrams but also opens new avenues for the rigorous characterisation of multicomponent phase-separating systems.

## I. INTRODUCTION

Biomolecular condensate formation in cells has emerged as a new paradigm in which nature achieves compartmentalisation of macromolecules — including proteins and RNAs — with thermodynamic principles [1, 2]. The relevance of these condensates in functions and diseases [3, 4] motivates experimental effort in understanding the driving forces [5–7] and molecular grammar of condensate formation [8, 9]. A commonly measured descriptor of phase separation is the phase diagram that indicates at which solute concentrations phase separation occurs, and the phase separation propensity is usually deduced by assessing how the phase diagram changes upon a modulation or change in polymer sequence [10, 11]. So far, phase diagrams have only been assessed qualitatively, and a general, quantitative framework is lacking that would allow one to systematically derive analytical relations for phase boundary shapes and to extract physical parameters from these measurements.

A major difficulty towards such a framework is the high-dimensional nature of the concentration space and associated phase boundary, which comprises a dense branch and a dilute branch connected by tie lines (Fig. 1a). In *in vitro* and *in cellulo* experiments, many solvent components, such as salt, buffer molecules, crowders, and pH, are present, and the measured phase diagrams usually represent low-dimensional experimental slices of the full concentration space (Fig. 1a). Mathematically, we denote by *c*^*α*^ the total molar concentration of solute *α*, where *α* = 1, 2, …, *N* for *N* solute species^1^. *The dilute and dense phase molar concentrations are denoted as* 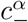 and 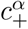 respectively. A typical 2-dimensional phase diagram measured along the *c*^1^ and *c*^2^ axes (without loss of generality) can be represented by a plane 𝒮_12_ intersecting the dilute phase branch, and the intersection is the measured phase boundary, equivalent to 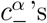 evaluated on 𝒮_12_ (Fig. 1a). A comprehensive theory would need to account for interactions among all *N* components and then solve the globally constrained optimisation problem to give the phase boundary, before a low-dimensional section of it can be fitted to experimental data. Although detailed theories have been built in the past to account for effects of pH [12, 13], crowders [14], and salt [15, 16], combining the effect of all possible components is analytically intractable. Model-free measurement approaches are emerging that allow relative energetics of phase-separating systems to be extracted in the presence of an arbitrary number of solutes using dilute phase measurements [6, 7, 17]. Can a similar method be developed to interpret phase diagrams without concentration measurements?

**FIG. 1.**
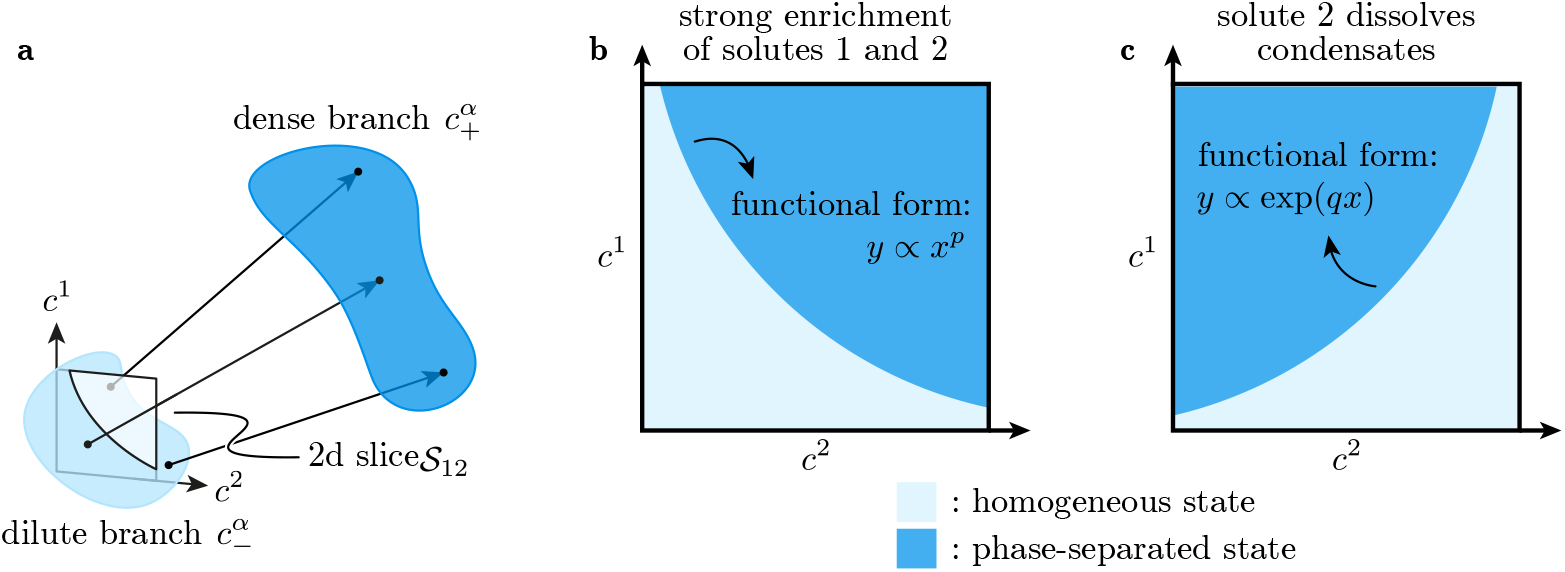
Phase boundary scaling relations for low-dimensional measurements of multicomponent systems. **a** In the concentration space with many solute components, the dilute 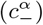 and dense 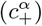 branches of the phase boundary are high-dimensional. Typical experiments only probe a small discrete set of components and hence measure a low-dimensional slice (𝒮_12_) of the dilute phase boundary. **b** When both solutes 1 and 2 are strongly enriched in the dense phase, possibly due to heterotypic attraction, a polynomial scaling relation is expected of the phase boundary, with the exponent related to their molar ratio in the dense phase. **c** If instead the solute 2 dissolves condensates comprising solute 1, the phase boundary has a positive gradient and an exponential functional form.

In this work, we derive an expression for the multi-component phase boundary normal vector, and evaluate it as a power series by assuming a generic free energy function encompassing translational entropy and many-body interactions. We study leading order contributions of this vector for the experimentally relevant 𝒮_12_ slice, which lead to analytical expressions of the phase boundary on 𝒮_12_. These give simple shapes in two scenarios: strong association, where entropic effects dominate for both components (Fig. 1b), and dissolution of condensates due to 3 or more-body interactions (Fig. 1c). These two cases capture the central effects of the two leading order contributions of the phase boundary normal expression, and higher-order interactions can be systematically incorporated. We make extensive use of the theoretical results to analyse phase boundary data for a wide range of experimental systems, with accompanying analysis scripts, and draw connections to mass action in chemical kinetics, the energy dominance framework, and solubility product for condensates. Taken together, our work establishes a new way of quantitative interpretation of low-dimensional phase boundary data.

## II. MAIN THEORETICAL RESULTS

The first scenario we explore is strong enrichment. For generic solutes 1 and 2, when their dense phase concentrations are much larger than their dilute phase concentrations, we show that the phase boundary on 𝒮_12_ satisfies a simple scaling relation (Fig. 1b, Appendix A)

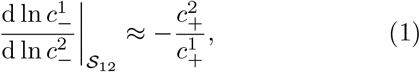

where 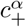 is the dense phase composition connected by a tie line to the point of evaluation on 𝒮_12_. This scaling leads to a polynomial functional form for the phase boundary on a *c*^1^ − *c*^2^ plot (Fig. 1b). Remarkably, this result is robust in the presence of other hidden components, and we verify this using numerically generated 2 and 3-component Flory-Huggins phase separation boundaries (Appendix B). This general scaling behaviour has a simple physical interpretation in terms of mass action when the activity of molecules in the dense phase is constant, a point we discuss later.

The second scenario we characterise is where increasing the concentration of a component leads to the dissolution of the dense phase, while the strong partitioning assumption of Eq. (1) only applies to one of the solutes.

Specifically, suppose component 1 is strongly enriched in the dense phase in the absence of solute 2, possibly driven by a heterotypic interaction with another solute 3 (without loss of generality). Furthermore, suppose addition of the solute 2 dissolves condensates, so the phase boundary in 𝒮_12_ has a positive gradient. We show that the phase boundary shape in 𝒮_12_ becomes an exponential function (Fig. 1c, Appendix A)

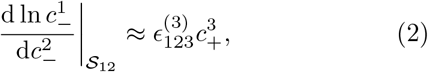

where 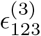 is a 3-body repulsion energy density involving solutes 1, 2, and 3, 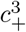 is the dense phase molar concentration of component 3, and 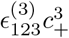 collectively represents a repulsion chemical potential for solute 2. If the solute 1 condensates are formed by homotypic interaction among solute 1 molecules, we can simply replace ‘3’ by ‘1’ on the right hand side of Eq. (2).

Comparing the left-hand sides of Eqs. (1) and (2), the main difference is changing d ln 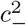 to 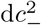. The logarithmic scaling originates from the entropic penalty of partitioning component 2 in associative phase separation, and 3-body disruptive interactions enter as a linear contribution to the phase boundary normal (Appendix A), leading to the difference in forms. Interestingly, a corollary of the theoretical results is that under the assumption of weak solvent entropy contributions, condensate dissolution by addition of a component can only arise from 3 or more-body interactions, and not possible with only 2-body interactions alone (Fig. 2, Appendix A). Dissolution observed in numerical computations of the 2-component Flory-Huggins model [18] arises from solvent entropy effects, and will be absent in the weak solvent limit. Typical molecular species that dissolve condensates are monovalent salts involved in charge screening [7, 19–21] and protein or RNA-binding partners that sequester condensate-forming proteins through complex formation [22, 23]. Our analysis suggests that these mechanisms should all broadly be considered as many-body interactions. This is sensible since these processes can be treated as 1 molecule disrupting attractions between 2 other molecules, and as such at least 3 molecules are involved in the dissolution mechanism (Fig. 2b). Many-body interactions can be a fruitful avenue for further investigations, as they are found to introduce multi-phase behaviours that are richer than binary interactions in numerical computations [24], and robust quantitative interpretation of 1-dimensional phase separation data also relies on 3-body interactions [25].

**FIG. 2.**
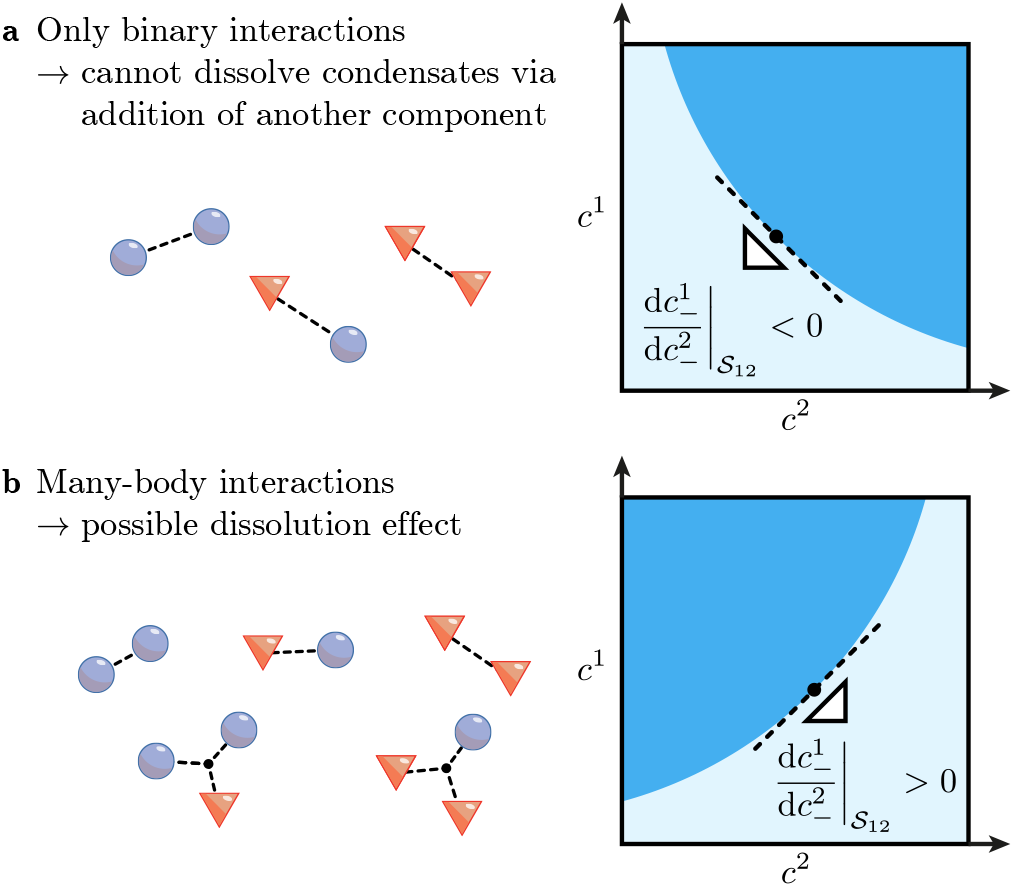
Order of interactions determines dissolution phenomenology. **a** When only binary interactions are present in the phase separation system, no condensate dissolution upon the addition of another component can be observed in the low-concentration limit where solvent entropy effects are negligible. **b** Dissolution is only possible in the presence of 3 or more-body interactions, which can be thought of as 1 molecule disrupting the attraction between 2 other molecules.

## III. APPLICATIONS TO EXPERIMENTAL DATA

To see how Eqs. (1) and (2) can be applied in practice, we analyse published as well as new data. We first take phase boundary data from [21] involving rotavirus proteins NSP2 and NSP5. Condensates formed by these proteins are key sites of viral replication [29]. To fit a power-law form to the phase boundary we plot the data in double logarithmic scale and use a support vector machine with linear kernel to estimate the phase boundary (Fig. 3a, 3b). Details of the analysis can be found on the GitHub repository [30]. This fitting gives a molar stoichiometry of 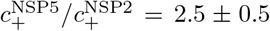. To validate the dense phase molar ratio, we perform a partitioning assay via direct concentration quantification by centrifuging a phase-separated sample followed by gel electrophoresis of the supernatant (Appendix C). In short, NSP2 and NSP5 are mixed at an equimolar concentration of 20 µM each at a final volume of 10 µl, giving a total of 0.2 nmol of each protein in the sample. The sample is centrifuged and the dilute phase is taken out as the supernatant. The amounts of proteins in the supernatant are quantified by comparing the gel stain intensities to samples containing only NSP5 or NSP2 (Fig. 3c). The result showed that (0.1988 ± 0.0003) nmol of NSP5 and (0.073 ± 0.006) nmol of NSP2 entered the dense phase, giving a dense phase molar ratio of 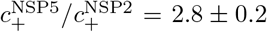, consistent with the phase boundary fitting result (Fig. 3d, 3e). Using the dense phase molar ratio, we can further give the energy dominance [7] ratio between these proteins. The dominance *D*^*α*^ is the fraction of free energy decrease due to solute *α* relative to the total free energy decrease, and satisfies 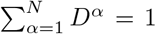 [7]. We show that for 2 generic components (Appendix A)

**FIG. 3.**
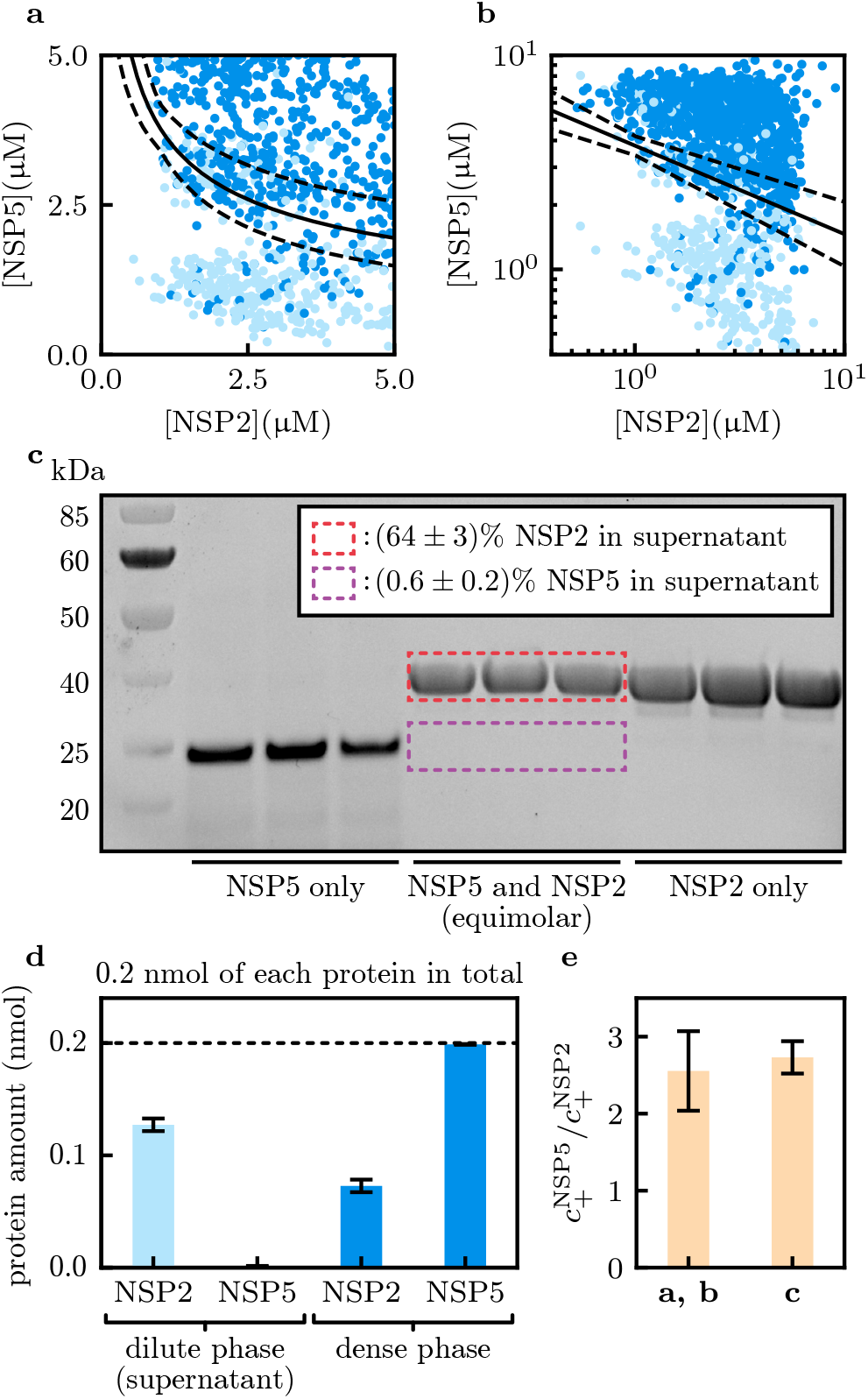
Phase boundary informs on the stoichiometry of viral protein condensates. Phase boundary data from [21] is reproduced here in **a** linear and **b** double logarithmic scales. Light and dark blue scatter points correspond to homogeneous and phase separated samples respectively. We analyse the boundary following the scaling relation Eq. (1). Black solid lines are power-law fits with errors in black dashed lines, giving a dense phase molar ratio of 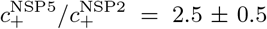. **c** Gel electrophoresis results for quantifying NSP5 and NSP2 dense phase molar ratio. Equal concentrations of NSP2 and NSP5 are mixed in solution to form condensates, and supernatants of samples with only NSP5 (left), NSP5 and NSP2 (middle), and only NSP2 (right) are measured. **d** Gel electrophoresis gave amount of proteins in the dilute phase (left, light blue), and the rest are in the dense phase (right, dark blue). This gives 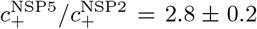. **e** Summary of dense phase molar ratios.

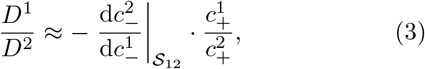

where 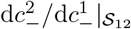 is the inverse of the phase boundary gradient. This implies that the phase boundary alone allows us to deduce the relative energetic contribution of solutes to phase separation, and a dominance ratio of *D*^NSP5^*/D*^NSP2^ ≈5 is estimated from data. This means the free energy decrease associated with NSP5 is about 5 times that associated with NSP2.

The NSP5/NSP2 condensate system is relevant *in cellulo* and *in vivo*, while many *in vitro* condensates are constituted using the macromolecule polyethylene glycol (PEG). PEG has been used by the condensate community as an inert crowder in the past but was later shown to have associative interactions in some cases [18], so we revisit the phase separation of the protein Fused in Sarcoma (FUS) with PEG (molecular weight 6000 dalton, or 6 kDa, in our experiments). We measured the FUS/PEG phase boundary (Appendix D) and found via fitting to Eq. (1) the dense phase molar ratio 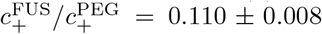 (Fig. 4a) [30]. In the past, this molar ratio has been estimated using the dilute phase measurement approach [18, 19], which gave 0.0061 ±0.0005 for PEG (20 kDa) and 0.0039 ± 0.0008 for PEG (10 kDa) in [18], significantly smaller than that measured here. We trace this inconsistency to a correction factor due to the measurement approach developed in [18]. In short, the approach outlined in [18] aims to find two compositions of total [FUS] and [PEG] that give the same dilute phase [FUS]. These points would have been on the same tie line if the system was entirely 2-dimensional, but this breaks down if more ‘hidden’ dimensions are present [7, 19]. A brief summary of the results from a more rigorous analysis [7] is given in Appendix E. Denoting by *ζ* the ratio between the FUS/PEG dense phase stoichiometry measured using the dilute phase approach and that of the true stoichiometry, we have

**FIG. 4.**
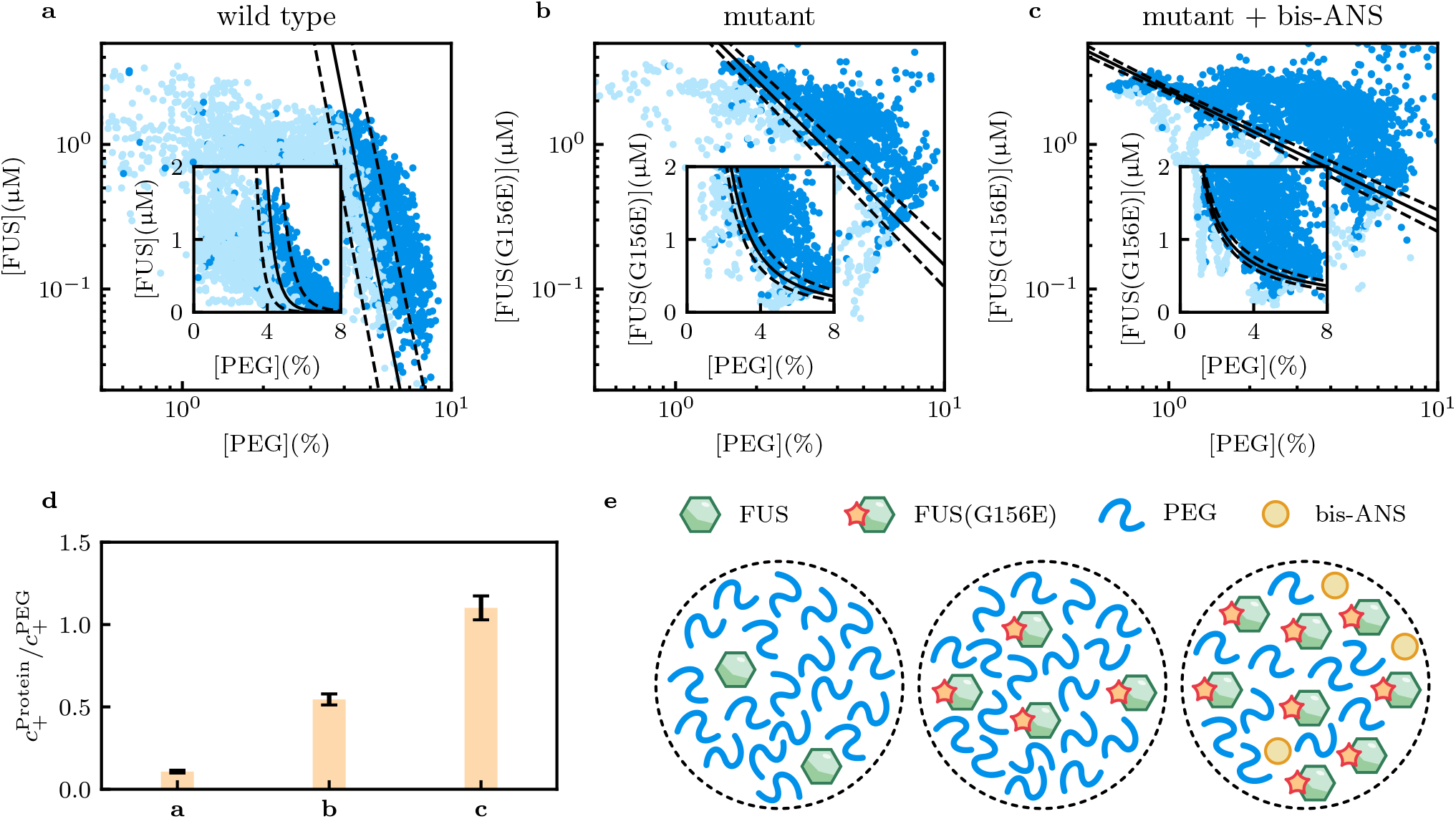
Phase behaviour of FUS with PEG. **a** PhaseScan experiment performed with wildtype FUS is analysed in double logarithmic scale, Giving a dense phase molar ratio 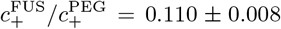. Inset: same data plotted in linear scale. **b, c** We take PhaseScan data of the mutant FUS(G156E) from [21] in the absence and presence of the small molecule modulator bis-ANS. The mutant FUS is more preferentially enriched than the wild type FUS, with a dense phase molar ratio 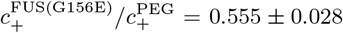. When the small molecule bis-ANS is added, this further increases to 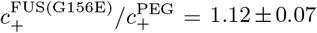. The phase boundary also shifts towards lower [PEG], indicating bis-ANS enhances the phase separation propensity of FUS(G156E) [21, 26]. **d** Summary of protein-to-PEG dense phase molar ratios. **e** Graphical illustration of the estimated condensate stoichiometry values.

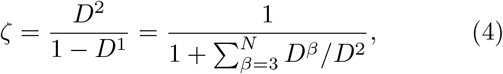

where FUS and PEG are solutes 1 and 2, and the sum is over all other solutes. We estimate *ζ* ≈ 0.02 ≪ 1 (Appendix E) by comparing results from [18] to the results obtained here, and deduce that *D*^2^ is small, meaning PEG is only contributing weakly to the system free energy decrease. This is further corroborated by estimating the FUS/PEG dominance ratio using Eq. (3). The phase boundary gradient is 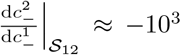 since the boundary varies in the µM range for FUS but mM range for PEG (1% in PEG corresponds to 1.6 mM in molar concentration), leading to 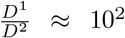. Using the measured FUS dominance *D*^1^ ≈ 0.6 in this concentration regime [7] we can estimate *D*^2^ ≈ 0.006, leading to 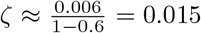, consistent with the earlier value.

Next, we demonstrate the utility of phase boundary scaling analysis in the context of modulation studies. In [21], phase diagrams between the mutant FUS(G156E) and PEG (molecular weight 6 kDa) are measured, in the presence or absence of the small molecule bis-ANS. Compared to the wild type FUS discussed before, the G156E mutant (where a glycine residue is replaced by a glutamic acid residue at position 156) is more prone to aggregation and implicated in the development of Amyotrophic Lateral Sclerosis (ALS) [26]. Repeating the log-log scaling analysis, we obtain for the dense phase FUS(G156E)-to-PEG molar ratio as 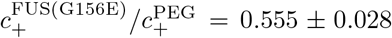 (Fig. 4b), significantly higher than that obtained for the wild type FUS (Fig. 4a). Upon addition of bis-ANS, which is documented to be a potent modulator of phase behaviour [31], this ratio further increases to 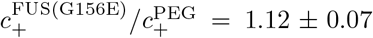(Fig. 4c) [30]. These results are in agreement with the fact that the G156E mutation enhances FUS self-association [26], and the role of bis-ANS as a bivalent linker that can strengthen hydrophobic interactions crucial for FUS condensate formation [32](Fig. 4d, 4e).

As a more biologically relevant example, we analyse data from [27] where the phase behaviour of the BNI1 RNA with protein Whi3 was studied. BNI1 RNA is an mRNA encoding the BNI1 protein, previously documented to phase-separate with Whi3 and can dissolve condensates at high RNA concentrations [28]. BNI1 RNA has 5 cognate binding sites for Whi3, and in [27] phase diagrams of the wildtype BNI1 RNA as well as a mutant RNA with cognate sites shuffled were measured (Fig. 5a, 5b). We fit the phase boundary in log-log scale, using only low-concentration data of RNA and Whi3 to avoid the dissolution branch, and observe that the RNA-to-Whi3 molar ratio decreases upon cognate site shuffling, from 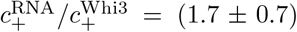 to 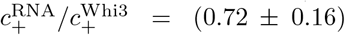 (Fig. 5b, 5c) [30]. The large fitting error for the wildtype RNA stems from the complex phase boundary shape, which shows dissolution of condensates by RNA as well as Whi3. Analysis of this dissolution branch can in principle be performed using Eq. (2), however the data points are sparse in this case. Instead, we use data from [28], where the dissolution effect of CLN3 RNA, an mRNA encoding the protein CLN3, is mapped out against Whi3.

**FIG. 5.**
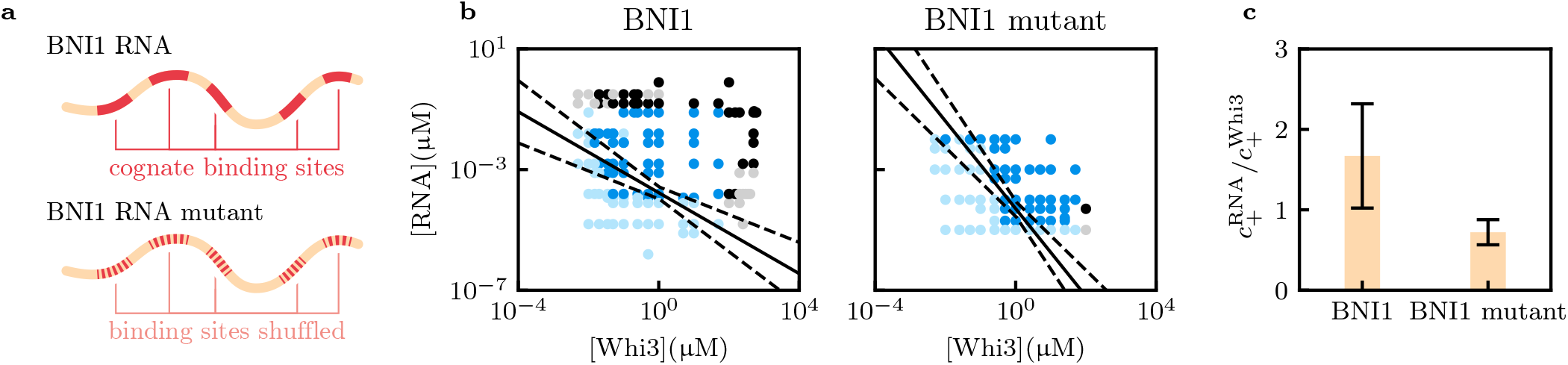
Whi3/RNA phase behaviour changes with RNA cognate binding sites shuffling. **a** Phase behaviour of BNI1 RNA and Whi3 were studied in [27]. BNI1 RNA (top) has 5 cognate binding sites for Whi3, and a mutant with the cognate sites shuffled was also studied (bottom). **b** Phase diagrams of Whi3 against the wildtype (left) and shuffled (right) BNI1 RNA measured in [27]. For phase boundary fitting, we only use points with [Whi3] *<* 100 µM and [RNA] *<* 0.1 µM, as points with higher concentrations might break the strong enrichment assumption and incur dissolution effects [28]. Light and dark blue scatter points are homogeneous and phase-separated data used for the fitting. Grey (homogeneous) and black (phase-separated) points are masked out during fitting. **c** Dense phase molar ratios of the RNAs against Whi3. A decrease in relative RNA content from 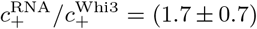 to 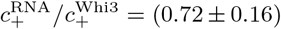 is observed here after shuffling the cognate sites, as expected.

Following the notation of Eq. (2), since RNA is dissolving condensates, we assign index 1 to Whi3 and index 2 to RNA. The Whi3/RNA phase boundary shape is then [compare to Eq. (2)]

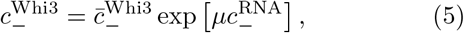

where the constant right-hand-side of Eq. (2) is collapsed into a single variable *µ* since we do not have prior information on which 3-body interaction is responsible for the dissolution. *µ* has the dimension of *k*_B_*T* (in this work we have set *k*_B_*T* = 1) per concentration unit and represents an effective repulsion chemical potential for RNA molecules. The inverse of *µ* then provides an RNA concentration scale over which dissolution occurs. We fit the phase boundary to a straight line in a log-linear plot, and fitting parameters are 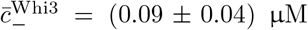 and *µ* = 9 *±* 3 *k*_B_*T/*µM, so the CNL3 RNA dissolution occurs over a characteristic range of ~0.1 µM. Detailed analysis scripts can be found in [30]. We note that Eq. (5) indeed captures the phase boundary shape to a good extent (Fig. 6a), however the sampling in [28] was done in double logarithmic scale, leading to data sparsity in linear scale (Fig. 6b). To further demonstrate the use of Eq. (2) we turn to [7], where dissolution of G3BP1 condensates by the small molecule suramin as well as poly(A)-RNA was studied [21].

**FIG. 6.**
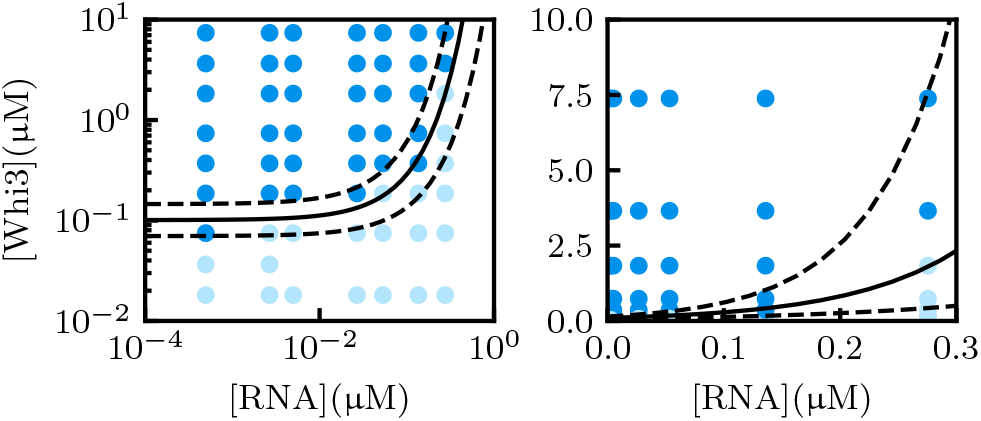
Exponential function captures CNL3 RNA/Whi3 dissolution boundary. We analyse phase separation data from [28] using Eq. (5), and plot the results in double logarithmic (left) and linear (right) scales. The original sampling was performed in double logarithmic scale, so data appears sparse in the linear scale. Fitting an exponential function leads to a characteristic CLN3 RNA chemical potential of *µ* = 9 *±* 3 *k*_B_*T/*µM.

G3BP1 is an RNA-binding protein central to stress granule formation [34]. In [7], phase separation of G3BP1 with poly(A)-RNA is studied under the dissolution effect of the small molecule suramin, which binds to G3BP1 and disrupts G3BP1/RNA interaction. Using data from [7], we plot the 2-dimensional G3BP1/suramin phase boundaries and fit exponential curves to them, at various levels of total [RNA] (Fig. 7a) [30]. The fitting function is [compare to Eq. (2)]

**FIG. 7.**
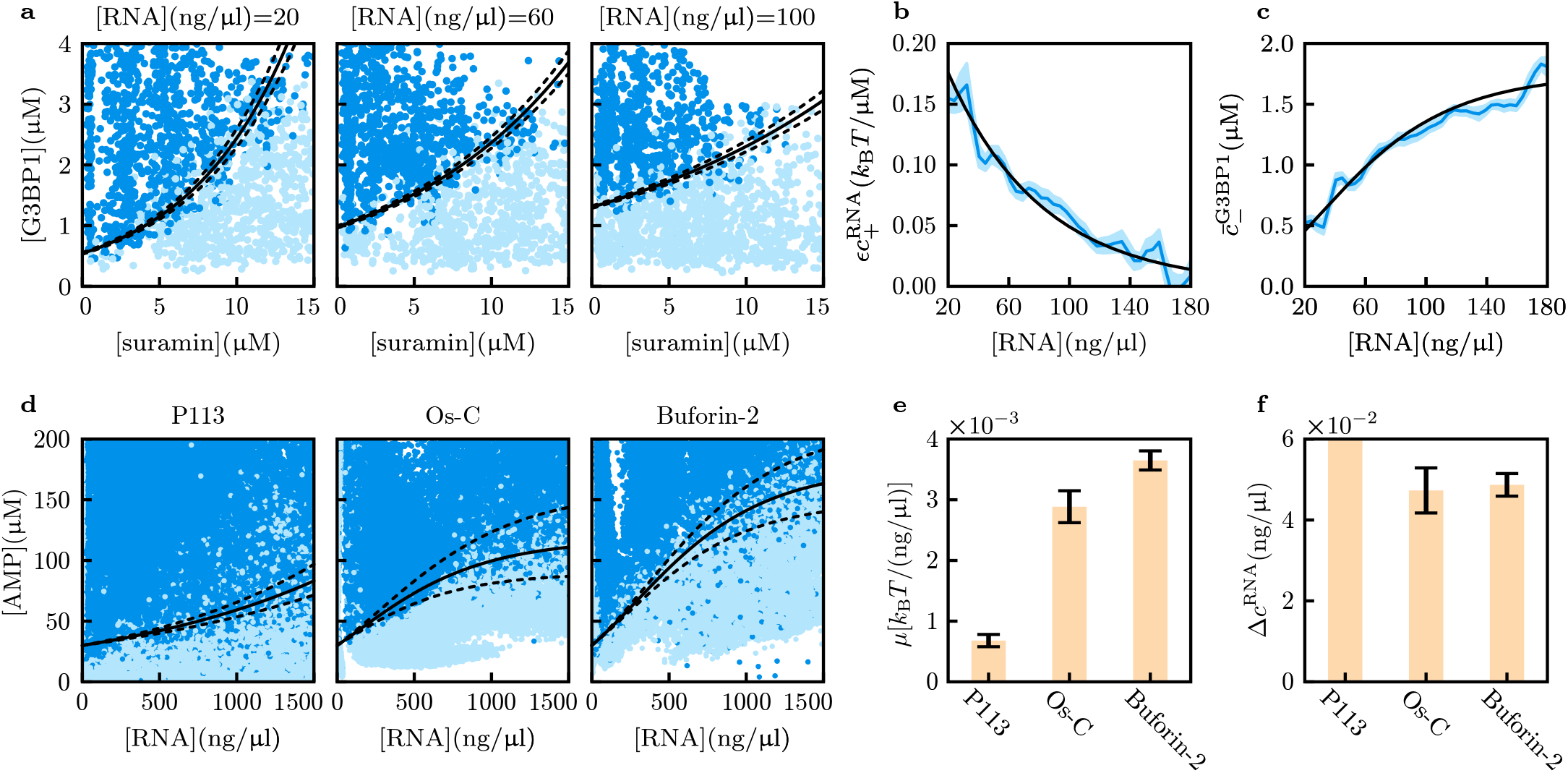
Condensate dissolution boundary analysis in more complex scenarios. **a** In [7], condensates formed by G3BP1 and poly(A)-RNA are dissolved by the small molecule suramin at varying levels of total [RNA]. An exponential functional form Eq. (6) is fitted to the phase boundary (black lines). **b** The fitting constant 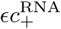 decreases as total [RNA] increases due to the dissolution effect of RNA. This implies the dense phase [RNA] decreases with total [RNA] (blue solid line, and blue shade represents the error estimate). Black solid line is an exponential fit using Eq. (7). **c** The G3BP1 saturation concentration in the absence of suramin increases with total [RNA]. This is effectively the G3BP1/RNA phase boundary, and the simple form of Eq. (2) cannot explain its concave shape. Instead, drawing from the hint that dense phase [RNA] is changing at different total [RNA], we use Eq. (8) instead (black solid line). **d** Phase boundaries of three anti-microbial peptides against poly(A)-RNA from [33]. Black solid and dashed lines are fits using Eq. (9). **e** The effective repulsion chemical potential *µ* is lowest for P113, also reflected in the least dissolution effect among the AMPs. Buforin-2 has the highest *µ*, in line with its strongest dissolution tendency. **f** Characteristic RNA concentration Δ*c*^RNA^ over which dense phase [RNA] decreases. The value obtained for P113 is very large, recovering the functional form of Eq. (2). Δ*c*^RNA^ estimated for Os-C and Buforin-2 are however similar, so the difference seen in experiments can be attributed to repulsion energy.

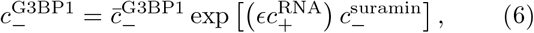

where 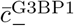 is the saturation concentration of G3BP1 in the absence of suramin, and the G3BP1/RNA/suramin 3-body repulsion energy is written as *ϵ* for brevity. Since *ϵ* is a constant, changes in 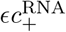 at different total [RNA] should arise from changes in dense phase [RNA], so we deduce that as RNA is added to the system, the dense phase [RNA] actually decreases (Fig. 7b). The fitted 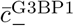 values are effectively the G3BP1/RNA phase boundary in the absence of suramin (Fig. 7c), which shows another dissolution trend. However, this boundary appears concave and cannot be rationalised using Eq. (2), so additional details need to be added to the theory to explain this phenomenon. In deriving Eq. (2), we assumed 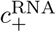 is constant, while Fig. 7b suggests that 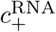 in fact decreases with 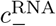. We thus propose a phenomenological functional form for 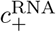 as a function of 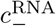 :

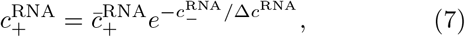

and find the fitting parameter Δ*c*^RNA^ = (33.7 ± 1.3) nM, or (70.8 ±2.8) ng/µL (black lines in Fig. 7b). This allows us to account for the change in dense phase RNA concentration when deriving the phase boundary scaling function of G3BP1 against RNA, leading to (Appendix F)

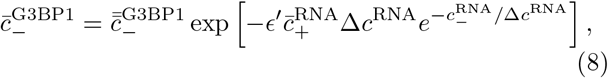

where 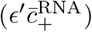 is a fitting parameter corresponding to the product of protein/RNA/RNA 3-body repulsion and dense phase RNA concentration at small RNA, 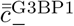 is another fitting parameter that fixes the *y*-intercept, and we take previously obtained Δ*c*^RNA^ value. Eq. (8) accounts for the data well (Fig. 7c), and this example illustrates the flexibility of the approach developed to analyse phase boundary scaling: by writing the phase boundary normal components as sums of terms, increasingly complex scenarios can be included.

Finally, we observe that the concave shape of Fig. 7c is rather common of protein/RNA dissolution phase boundaries, and hypothesise that despite the phenomenological nature of Eq. (7), the derived form Eq. (8) can still be used on other protein/RNA systems. We take phase boundary data from [33] where the phase behaviours of three anti-microbial peptides (AMPs), P113, Os-C, and Buforin-2, in the presence of poly(A)-RNA, were mapped out. Eq. (8) represents a 3-parameter fit if Δ*c*^RNA^ is not estimated, leading to a large fitting error. To better constrain the fitting, we use the following functional form

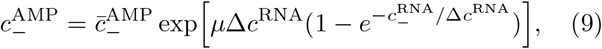

and fix 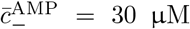 for all three datasets. This constrains the *y*-intercepts of the fits, and the fitting parameters are *µ*, which has the dimension of chemical potential for the repulsion energy, and Δ*c*^RNA^, a characteristic RNA concentration scale over which dissolution affects dense phase RNA concentration. Eq. (9) provides a good fit to the three phase boundaries (Fig. 7d), and detailed analysis code can be found in [30]. It was originally found in [33] that P113 has the highest phase-separation propensity with RNA, with the largest phase-separated region among the three AMPs (Fig. 7d). Indeed, comparing the fitted *µ* values shows that P113 has the smallest repulsive chemical potential, followed by Os-C and Buforin-2 (Fig. 7e). The Δ*c*^RNA^ estimated for P113 is very large, hinting that dense phase RNA concentration is hardly changing (Fig. 7f). Interestingly, Os-C and Buforin-2 have similar Δ*c*^RNA^, and the difference in measured phase boundary solely originates from the effective potential *µ* (Fig. 7f). In addition, since the RNA used is the same across these experiments, the repulsive term has to involve the AMP, further supporting the assumption used to derive Eq. (8) in Appendix F. Taken together, we conclude that Eqs. (2) and (9) can be helpful tools to extract quantitative information from experimentally measured phase boundaries when dissolution is observed.

## IV. DISCUSSION

Obtaining high-dimensional phase boundaries in theoretical models is a daunting task, and the framework we developed here represents an alternative route towards quantitative interpretation of experimental data. The two general phase behaviours we explored are observed in many multicomponent systems, and connections can be made with other existing theoretical frameworks too.

It is instructive to compare Eq. (1) with mass action in chemical kinetics. For a process where molecular species *A* and *B* combine to yield *C*

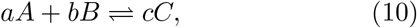

with *a, b, c* being the stoichiometric coefficients, we have at equilibrium *K*^eq^[*A*]^*a*^[*B*]^*b*^ = [*C*]^*c*^ for an equilibrium constant *K*^eq^. The derivation of this relation is mathematically similar to the binodal equations, where the chemical potentials of the *A* and *B* molecules in the monomeric state and the combined *C* state are equalised, originiating from global free energy minimisation with mass conservation constraints. Taking the logarithm of both sides of Eq. (10) and perturbing the concentrations, while assuming the concentration [*C*] is approximately a constant (such as *C* being in a different phase), we obtain

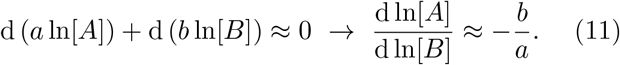

As such, Eq. (1) has a simple physical interpretation as a solubility product. In [35, 36], a similar solubility product idea is proposed, where the dilute phase solute concentrations are multiplied together instead of using the phase boundary values. Using the dilute phase response relations derived in [7] we show that the solubility product of [35, 36] is closely related to the phase boundary scaling Eq. (1) via dominance values of the two solutes as well as tie line vector components (Appendix G).

For condensate dissolution, we show that 3-body repulsion is necessary in practical situations and a generic exponential phase boundary shape ensues. The fitting parameter in this case relate to the 3-body repulsion energy and dense phase concentrations, so explicit interaction energy values could in principle be obtained if the dense phase concentration can be measured. Furthermore, we find that the concave phase boundary shape typically exhibited by protein/RNA systems can be rationalised by a changing dense phase RNA concentration, which we first deduced from the G3BP1/RNA/suramin dataset of [7] and subsequently deployed to analyse datasets of [33]. Experimental confirmation of Eq. (7) via dense phase quantification in the future will provide further support for the theory. Perhaps a more interesting direction this opens up is in linking the dissolution phenomenon with more detailed mechanisms, such as molecular binding. It is usually surmised that strong protein/RNA binding underlies stable complex formation and subsequent condensate dissolution, and modelling the binding as introducing a new inert molecular species could lead to a binding-associated free energy contribution as a series expansion at low concentrations, drawing connection between 3-body repulsion energy and the binding constant.

In sum, the general mathematical formulation of the phase boundary normal expression allows quantitative interpretation of phase diagrams without explicitly solving for the phase equilibrium. This work makes extensive use of the theoretical results to demonstrate applicability of the phase boundary expressions, and the accompanying datasets and analysis scripts can be valuable for future experimental investigations.

## Data and code availability

Curated datasets and analysis scripts can be found on https://github.com/dq219/phase-boundary-scaling.

## Conflict of interest

The authors declare no conflict of interest.

## Acknowledgement

This work has received support from the European Research Council under the Horizon 2020 research and innovation program (agreement ID 101001615), Transition Bio Limited. (D.Q. and R.M.S.), the EPSRC SBS DTP [2597129] (J.A.), and the Wellcome Trust [213437/Z/18/Z] (A.B.). The authors thank Dr. Timothy J. Welsh, Dr. William E. Arter, and Dr. Hannes Ausserwoger for helpful discussions.

## Appendix A Mathematical derivations

We adopt notations from [7]. In an *N*-component phase separation system, we denote the average volume fraction of component *α* as *ϕ*^*α*^, where *α* = 1, 2, …, *N*, and the free energy of a homogeneous composition *ϕ*^*α*^ as *f* (*ϕ*), where the superscript is dropped in function arguments for brevity. The unit thermal energy is set to unity *k*_B_*T* = 1. Given dilute and dense phase component volume fractions 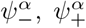, as well as the tie line vector connecting the points 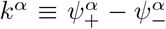, the chemical and osmotic equilibrium conditions are [7]

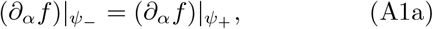

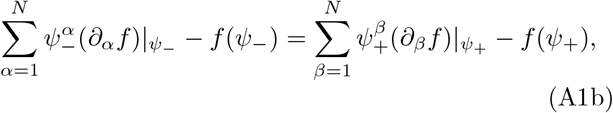

where 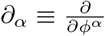. Perturbations of Eqs. (A1) lead to an expression of the phase boundary normal vector *n*_*α*_ [7], and we outline the detailed derivation here which was lacking before. Perturbing the left-hand-side of Eq. (A1a) gives

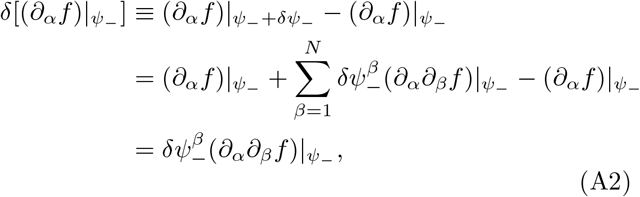

and repeating it for the right-hand-side dense phase point leads to

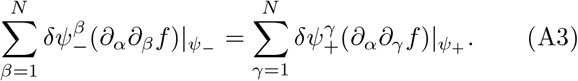

Similarly, we apply a perturbation to Eq. (A1b). We use the relations

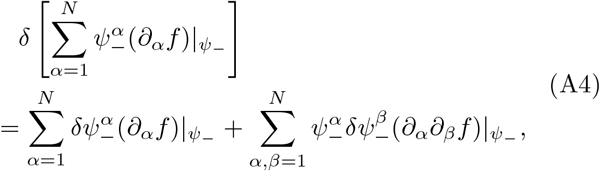

as well as

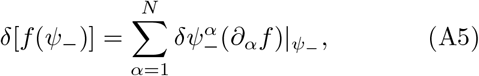

so that the first-order perturbation of Eq. (A1b) is

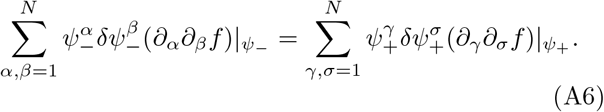

Substituting Eq. (A3) into the right-hand-side of Eq. (A6) gives

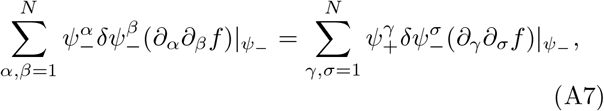

and using 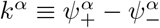 this can be simplified into

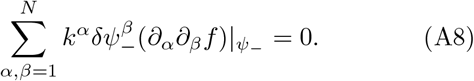

Since the phase boundary normal is defined by

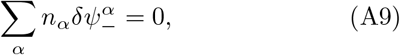

in essence, any perturbation must lie on this boundary and thus have zero overlap with the normal to the surface, direct inspection gives

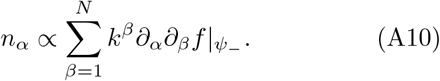

Eq. (A10) can in principle be used as it is, and one could further apply chemical potential equilibrium to simplify expressions. We now derive another form of *n*_*α*_ by combining Eq. (A10) with Eq. (A1a). Chemical potentials are defined as *µ*_*α*_(*ϕ*) ≡ (*∂*_*α*_*f*)|_*ϕ*_, and chemical equilibrium requires 0 = *µ*_*α*_(*ψ*_+_) − *µ*_*α*_(*ψ*_−_). In addition, we have 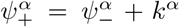 by definition, so that we can write chemical equilibrium in terms of quantities evaluated at the dilute phase 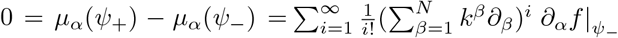. The *i* = 1 term is proportional to *n*_*α*_, so

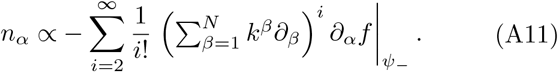

We now take as *f* (*ϕ*) a sum of components’ translational entropy, solvent entropy, and many-body interactions. It is convenient to define the solvent volume fraction as a function 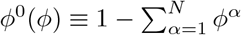, giving

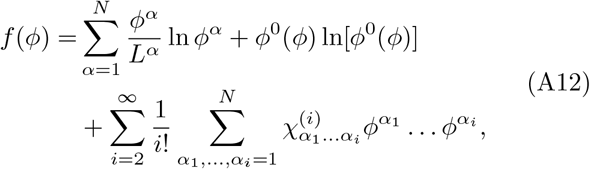

where *L*^*α*^ is the size of component *α*, and 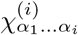 is the *i*-body contact energy. When only binary interactions 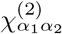 are considered we recover the Flory-Huggins model [37, 38]. Substituting Eq. (A12) into Eq. (A11) term by term leads to an expression for *n*_*α*_. The contribution to *n*_*α*_ from the first term of Eq. (A12) is

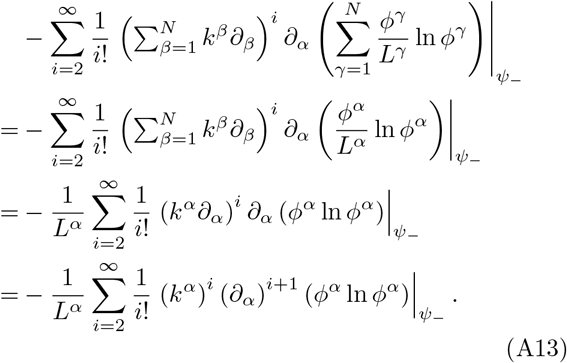

Note the generic relations

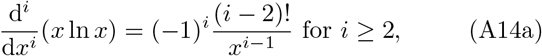

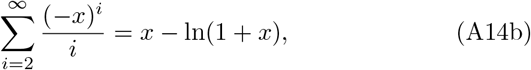

we can further simplify Eq. (A13) to

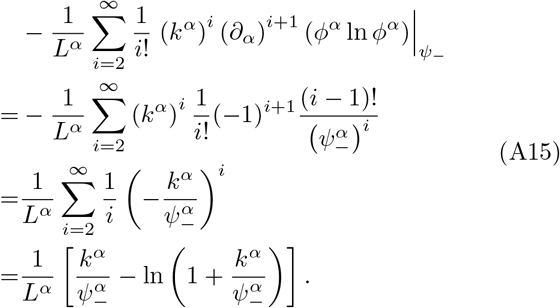

In the first line we used Eq. (A14a), and in the last line Eq. (A14b). The contribution from the entropic term of Eq. (A12) can be calculated in a similar fashion, leading to a compact form for *n*_*α*_:

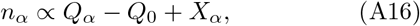

where

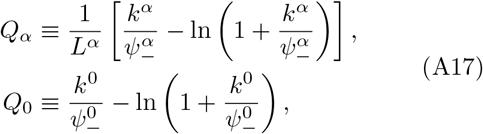

are terms arising from the entropic free energies, with shorthands 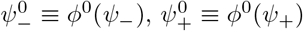, and 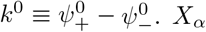 contains interaction terms and only interactions involving 3 and more bodies enter the *X*_*α*_ expression, with the leading term 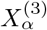

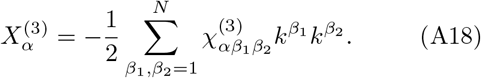

2-body interactions have disappeared as they are absorbed into *Q*_*α*_ through the chemical equilibrium. Notice all *Q*_*α*_’s and *Q*_0_ are non-negative since *x* − ln(1 + *x*) ≥ 0 ∀ *x* ∈ ℝ. We expect the solvent entropy contribution to be negligible in practical cases, *Q*_0_ ≈ 0. This is because the volume fraction of water is large in both dilute and dense phases in experiments, far from the logarithmic singularity, and furthermore a large *Q*_0_ would mean any molecule added to *in vitro* experiments can dissolve condensates, which is not the case. As a result, in the absence of 3 or more-body interactions (*X*_*α*_ = 0), the ratio between any two phase boundary normal components is always positive, and one cannot dissolve condensates via addition of another molecular species.

Eq. (A16) gives the general phase boundary normal direction. To make use of *n*_*α*_, note that for a vector 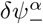 lying on the phase boundary, we have by definition

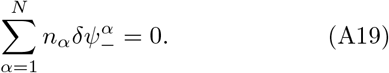

Since total concentrations of only components 1 and 2 are allowed to vary on 𝒮_12_, 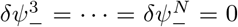. As a result,

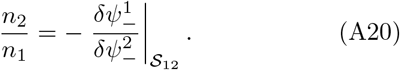

The right-hand side is closely related to the phase boundary gradient in 𝒮_12_, so an expression for *n*_2_*/n*_1_ allows a link between the experimental observable and the phase space to be established.

Assume solute 1 (without loss of generality) is strongly enriched such that 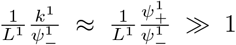. Meanwhile, since 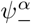 and *k*^*α*^ are of 𝒪 (1) (volume fractions), and the interaction *χ*’s are of 𝒪 (1) in units of thermal energy *k*_B_*T*, we have *X*_*α*_ ~ 𝒪 (1). In the strong enrichment regime, we can approximate

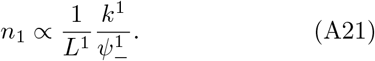

If there is a component 2 that is also strongly enriched, using Eq. (A20) we get

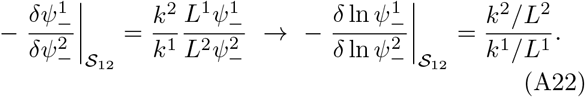

Finally, we convert volume fractions 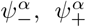 to their counterparts in molar concentration unit 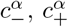 using

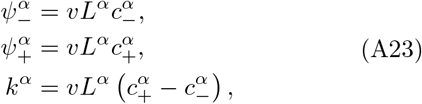

where *v* is the volume of one mole of lattice sites, to give

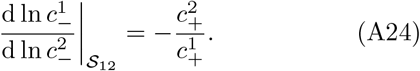

Here we also used 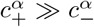. This is Eq. (1).

To give the dominance ratio between solutes 1 and 2, recall that [7]

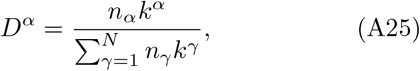

so

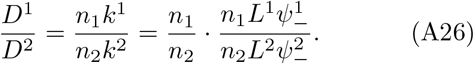

After algebraic simplifications we obtain

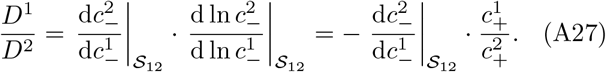

Next, we derive the phase boundary shape in the case of condensate dissolution. Suppose condensates formed by component 1 are dissolved by the addition of a component 2, through a ternary interaction with component 3 with 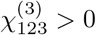 (or with component 1, in which case the derivation is similar). We use Eq. (A21) for *n*_1_ and take *n*_2_ ≈ *X*_2_, assuming component 2 partitioning and solvent entropy are weak factors. This allows us to understand the effect of 3-body interactions, and deviations of the phase boundary from the analytical form can be used to detect additional factors, such as a non-zero *Q*_2_ term. Adopting the simplifying assumptions leads to

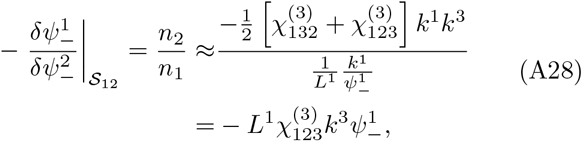

taking 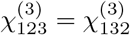. Re-arranging, this is

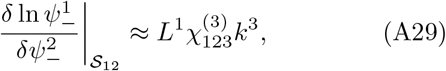

which gives an exponential form for the phase boundary in 𝒮_12_. In molar concentration units this gives

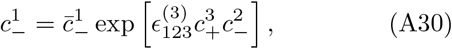

where 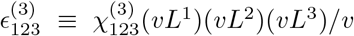 is the moleculelevel ternary interaction contact energy density in Eq. (2), and 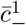 is the component 1 saturation concentration in the absence of component 2.

## Appendix B Scaling in a simple computational model

We verify the polynomial scaling of the dilute phase boundary by numerically computing the full phase diagrams of 2 and 3-component Flory-Huggins models. The free energy is taken to be, for *N* = 2 or *N* = 3:

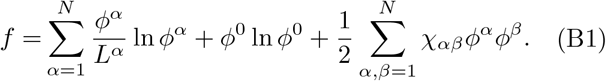

For the 2-component case, we set *L*^1^ = 40, *L*^2^ = 70, *χ*_11_ = −0.8, *χ*_12_ = *χ*_21_ = −3.0, and *χ*_22_ = −0.2. This choice of parameters results in strong enrichment of both components in the dense phase, while maintaining the solvent volume fractions to be of order 1 on the phase boundary (Fig. 8a, 8b). On a log-log plot, the dilute phase boundary is a straight line in the small-*ϕ* regime, and we fitted 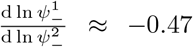. This gives 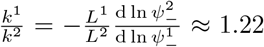, which is a good approximation since the exact value computed for points in the range 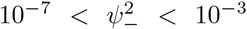 is 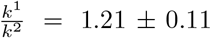. For the 3-component case, we keep the same parameters components 1 and 2, and set *L*^3^ = 5, *χ*_13_ = *χ*_31_ = 0.1, and *χ*_23_ = *χ*_32_ = 0.1. The tie lines are 3-dimensional vectors (Fig. 8c). The binodal boundary is computed using the convex hull algorithm and we take cross-sections of it at approximately constant *ϕ*^3^’s, using a window of Δ*ϕ*^3^ = 0.02. For each section, the fit the *ϕ*^1^ − *ϕ*^2^ boundary on a double logarithmic plot as before and estimate the *k*^1^*/k*^2^ ratio from the scaling exponent. Comparing this with *k*^1^*/k*^2^ explicitly computed using numerical data, we observe good agreement between the two methods (Fig. 8d).

**FIG. 8.**
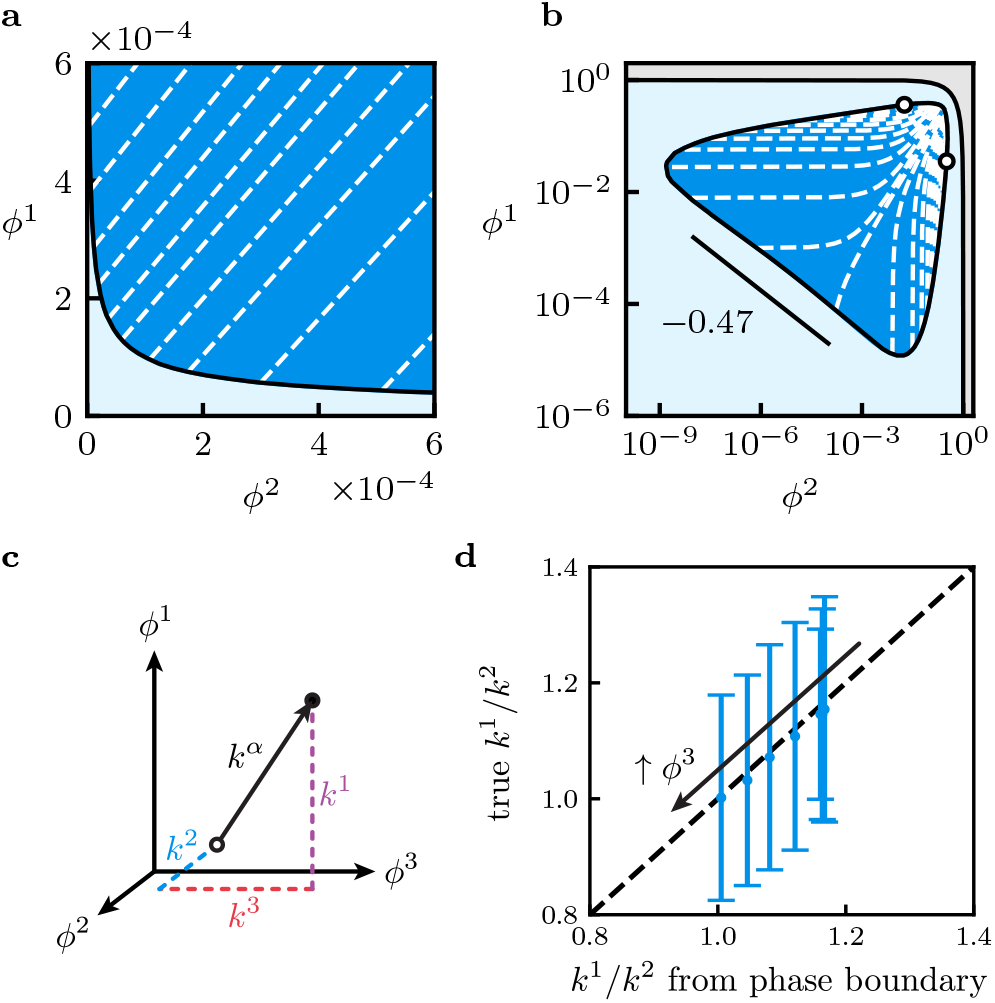
Numerical investigations using the Flory-Huggins model. **a, b** 2-component phase diagrams in linear and log scales. Light blue regions are homogeneous, and dark blue regions are phase-separated. White dashed lines are tie lines and hollow markers are critical points. The grey region is physically forbidden. Polynomial scaling of the phase boundary is observed at small *ϕ*^1^ and *ϕ*^2^, with the exponent relating to the tie line component ratio. **c** In the 3-component system, each tie line is a 3-dimensional vector. **d** We take slices of constant *ϕ*^3^ in phase space and compute *k*^1^*/k*^2^ from numerical data as well as the double logarithmic fitting, and they agree well with each other.

## Appendix C Experimental details for NSP5/NSP2 dense phase quantification

Expression of NSP5 nStrep (strain RF) and purification/solubilisation of inclusion bodies was conducted as described in [39, 40]. Solubilised inclusion bodies were dialysed [40], and the cleared supernatant was further purified using a Strep-tactin superflow column (iba lifescience). The column was washed with 1 M NaCl in 20 mM MOPS buffer at pH 7.1, and NSP5 was eluted using a 20 mM MOPS buffer containing 1 M NaCl and 2.5 mM Desthiobiotin (Sigma Aldrich) at pH 7.1. Protein-containing fractions were concentrated using an Amicon Spin column (Millipore) with a molecular weight cut-off of 10 kDa, and a final NaCl concentration of 150 mM was adjusted by addition of an appropriate volume of 50 mM MOPS at pH 7.1. Recombinant NSP2 cHis (strain RF) was expressed and purified as described in [29, 39]. Protein concentration and purity was monitored using a Spectrophotometer (Extinction coefficient NSP5: 10555 and NSP2: 41620) and SDS-PAGE, respectively.

To perform the condensate partitioning assay, NSP2 and NSP5 were mixed at an equimolar concentration to form a final sample of 10 µl in volume, with 20 µM of each protein. The sample is centrifuged for 30 minutes at 16000 rcf, alongside controls containing only NSP5 or NSP2. The supernatant was mixed with 4x mPAGE LDS Sample Buffer (Merck) and boiled at 70^°^ C for 10 minutes. A 12% mPAGE Bis-Tris Polyacrylamide Precast Gels (Merck) was run in MES buffer for sample visualisation, the BenchMark Pre-stained Protein Ladder (Thermo Fisher Scientific) was used as a protein standard. The gel was stained in Bio-Safe Coomassie Stain (Bio-Rad Laboratories) and imaged using a ChemiDoc imaging system (Bio-Rad Laboratories). The intensities and volumes of individual protein bands were quantified using the Image Lab software v.6.1.0 (Bio-Rad Laboratories). The intensity values of the NSP5 and NSP2-only samples were used as a standard to estimate the remaining protein concentration in the supernatant of the mixed NSP5/NSP2 sample.

## Appendix D Experimental conditions for FUS/PEG PhaseScan

For the PhaseScan [21] experiment we purify FUS protein using a previously established protocol [41]. PEG with molecular weight 6000 dalton is purchased from Sigma, and 5 µM of Alexa647 dye is added to the PEG inlet to track its concentration in droplets. Both FUS and PEG are dissolved in 50 mM Tris buffer with 150 mM KCl at pH 7.4.

## Appendix E Estimating dense phase molar ratio using dilute phase response

Using notations from Appendix A, the change in the dilute phase of a sample 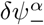 in response to a change in total composition *δϕ*^*α*^ (not constrained on 𝒮_12_) in the vicinity of the phase boundary is given by

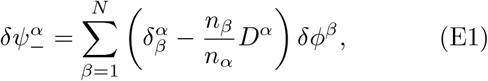

so that when *δϕ*^*α*^ is constrained on 𝒮_12_ we have an expression for the dilute phase component 1 concentration

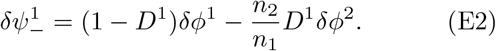

Setting 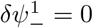 then gives a set of total compositions on 𝒮_12_ with a fixed dilute phase component 1 concentration, and this is

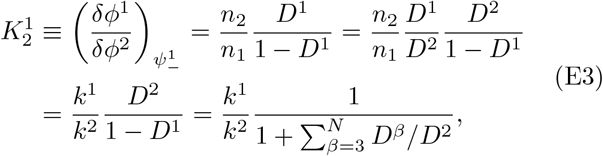

so the correction factor *ζ* is

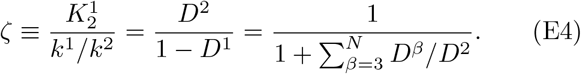

To estimate *ζ* using the molar ratio obtained from the scaling and from the dilute phase measurement approach, we have to take into account the different molecular weights of PEG used in these experiments. The 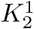 obtained in molar units are 0.0061 ± 0.0005 for PEG (20 kDa) and 0.0039 ± 0.0008 for PEG (10 kDa) [18], which appeared to be proportional to the molecular weight of PEG used (very crudely). For PEG of 6 kDa, we thus estimate a 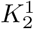 of 0.002 (in molar units), which is combined with the measured 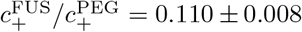 to give *ζ* ≈ 0.02.

## Appendix F Analysing G3BP1/RNA phase boundary

The dissolution branch of the G3BP1/RNA phase boundary (Fig. 7c) is concave, so Eq. (2) cannot be directly applied. Using indices 1 and 2 to denote G3BP1 and RNA, the problem lies in the assumption of a constant *k*^2^. The 3-dimensional PhaseScan data suggests *k*^2^ is in fact changing as a function of total RNA concentration (Fig. 7b), so the derivation leading to Eq. (A30) needs to be modified. From the data, we postulate that *k*^2^ changes with 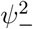 according to the functional form

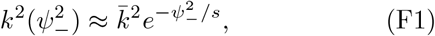

with constants 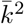 and *s*. The task now is to re-derive Eq. (A28). Since the phase boundary Fig. 7c corresponds to a system without suramin, we assume the dominant *n*_2_ contribution to be of the form

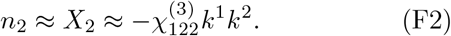

It is possible to add 3-body interaction terms corresponding to disruption of G3BP1/G3BP1 interaction by RNA 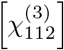 or disruption of RNA/RNA interaction 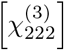. However, from the dilute phase G3BP1 measurement, it was postulated that RNA acts on interactions involving RNA [7], so we drop the 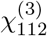 term. Furthermore, since we are assuming strong G3BP1 enrichment, we expect a large *k*^1^ and 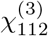 should dominate the *n*_2_ expression. We can now calculate

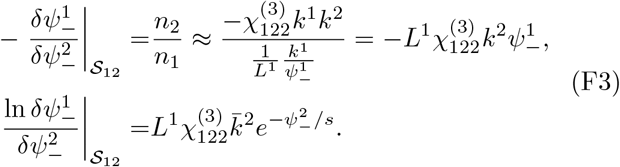

Integrating, this leads to

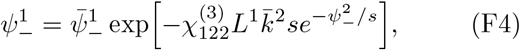

or, in molar units,

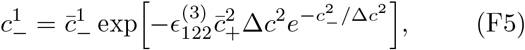

where 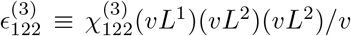 and Δ*c*^2^≡ υ *L*^2^*s*. To perform the fitting, first note that 1 ng/µl of the RNA corresponds to approximately 0.47 nM. Fitting Fig. 7b data to an exponential form yields Δ*c*^2^ ≈ (33.7 *±* 1.3) nM, or (70.8 *±* 2.8) ng/µl. Using this value of Δ*c*^2^ in Eq. (F3) we find 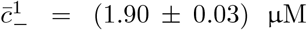, and 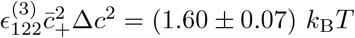 *k*_B_*T*.

## Appendix G Solubility product in the proximity of the phase boundary

On every point on the 2-dimensional slice of the phase boundary, we have complete information on the dilute composition since it is exactly the total composition in the limit of zero dense phase volume. However, for samples within the phase-separated regime, the dilute phase composition differs quickly from the total composition as the dilute phase point moves away from the 2-dimensional slice and into the *N*-dimensional space (Figure 1 a). As a result, multiplying the dilute phase concentrations within the phase-separated regime, as outlined in [35, 36], needs a separate treatment. We first take a point on the phase boundary with composition 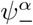 and bring it into the phase-separated regime by addition of some amounts of solutes 1 and 2, with this displacement denoted as *δϕ*^*α*^. The dilute phase composition of this new sample is 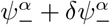 where 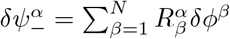 and 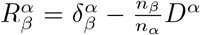 near the phase boundary, with 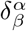 the Kronecker delta [7]. Direct substitution gives dilute compositions of components 1 and 2 as

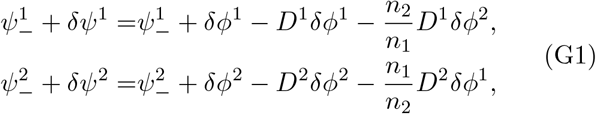

and 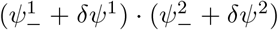 is effectively the solubility product referred to in [35, 36, 42]. The present framework however implies the appropriate product combination on the phase boundary should have the concentrations raised to the power of their dense phase molar compositions. We thus define the product *P* as

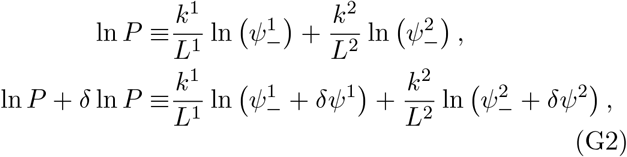

so that *P* is a constant on the phase boundary in 𝒮_12_. Treating *δϕ*^1,2^ as small perturbations, direct substitution leads to

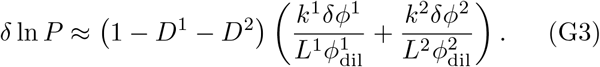

In the case where *D*^1^ + *D*^2^ = 1, we have *δ* ln *P* = 0, and the system is effectively 2-dimensional. On the other hand, if *D*^1^ = *D*^2^ = 0, ln *P* +*δ* ln *P* equals to the product formed using the total concentrations of 1 and 2 instead of the dilute concentrations, which is the product in the absence of phase separation.

## Footnotes

1 The superscript label follows conventions from differential geometry. For a vector quantity, a superscript *α* carries unit of concentration of *α*, while a subscript carries the inverse concentration instead.

## Notes

https://github.com/dq219/phase-boundary-scaling

### Competing Interest Statement

The authors have declared no competing interest.

